# "Smart Tanks": An Intelligent High-Throughput Intervention Testing Platform in *Daphnia*

**DOI:** 10.1101/2021.05.30.446339

**Authors:** Yongmin Cho, Rachael A. Jonas-Closs, Lev Y. Yampolsky, Marc W. Kirschner, Leonid Peshkin

## Abstract

We present a novel platform for testing the effect of interventions on life- and health-span of a short-lived semi transparent freshwater organism, sensitive to drugs with complex behavior and physiology – the planktonic crustacean *Daphnia magna*. Within this platform, dozens of complex behavioural features of both routine motion and response to stimuli are continuously accurately quantified for large homogeneous cohorts via an automated phenotyping pipeline. We build predictive machine learning models calibrated using chronological age and extrapolate onto phenotypic age. We further apply the model to estimate the phenotypic age under pharmacological perturbation. Our platform provides a scalable framework for drug screening and characterization in both life-long and instant assays as illustrated using long term dose response profile of metformin and short term assay of such well-studied substances as caffeine and alcohol.

## Introduction

Developing pharmacological interventions that slow down the aging process and consequently postpone the onset and progression of age-associated diseases is highly sought after. The aging process can be actively regulated by multiple interventions such as environmental, genetic, and pharmacological factors. Specifically, a large body of research has found that a reduction in caloric intake can extend lifespan and delay disease onset in a wide range of species from nematodes to primates^1-5^. A variety of heritable variants have been identified that to some extent mimic caloric restriction. For example, a *Caenorhabditis elegans* mutant for the insulin receptor *daf-2* can live two to three times longer than wild-type animals^6^, and flies and mice that have mutations in the insulin or insulin-like growth factor-1 receptor gene similarly show an enhanced lifespan^7-9^. Drugs that affect hypothesized metabolic pathways responsible for caloric restriction, such as rapamycin, metformin, and resveratrol are being studied for their potential to enhance lifespan in several organisms from nematodes to mice^10-13^. Pharmacological interventions are currently the most practical strategy for affecting aging in humans, avoiding the technical and ethical problems with genetic interventions and the difficulty of maintaining an unpleasant, life-long calorie-restricted diet. However, since the mechanisms driving the aging process are not well understood, there currently exist few druggable targets for anti-aging treatments and therefore the evaluation of drug effects on the aging process requires the development of new high-throughput screening platforms.

Quantitative biomarkers of aging are critical to developing novel therapeutic approaches. Since individuals may not age at the same rate, quantitative biomarkers of aging are valuable tools to measure physiological age, assess the extent of healthy aging, and potentially predict not only health and lifespan but also age-related outcomes for individuals within a population, even at the early age. Molecular biomarkers (often based on gene expression) are robust quantitative metrics but often require sacrifice of the subject, laborious sample processing, and constitute a single data end-point. Phenotypic biomarkers can be harder to quantify but are fairly easy to obtain, non-invasive, and therefore are possible to repeatedly assay over the entire life of the subject and in future generations, whereas molecular ones can reflect some of the molecular mechanisms underlying the aging process^14,15^. In this regard, walking speed, the ‘chair stand test’, standing balance, and body mass index are well-known biomarkers of aging in humans^14^. While these assays demonstrate the possibility of using phenotypic information as aging biomarkers, for obvious ethical and time considerations humans are an unsuitable model for the initial screening of chemical compounds or other interventions that may ameliorate age-associated phenotypic declines. Therefore, the development of novel model systems and their aging biomarkers will enable the discovery of potential anti-aging therapeutic strategies for humans.

A study of aging in a simple, short-lived model organism is extremely attractive. The small crustacean *Daphnia* (also called “water flea”) promises to be a powerful pharmacological model organism for several reasons: 1) it is a diploid, parthenogenetic species with a relatively short median lifespan (50 ∼ 100 days depending on environmental conditions)^16-19^, 2) it has a short reproductive cycle (e.g., a female can produce a clutch of 1-25 neonates every instar^20^, and 3) its genome and complex body plan are significantly more homologous to humans than a common aging model worm *C. elegans*, thus allowing more human-relevant, tissue-specific manifestations of aging to be analyzed^21^. These properties of *Daphnia* allow short timeline experiments with a high sample size for aging research, while their complex phenotypes provide opportunities to build strong phenotype-based biomarkers to assay the effects of drugs on the aging process. One of the most important parameters in drug development is the absorption, distribution, metabolism, excretion, and toxicity (ADME-Tox) of drugs. *Daphnia* allow the ease of perturbation by small molecules compared to other short-lived invertebrate model organisms such as *C. elegans* and *Drosophila*, which have impermeable cuticles that form strong barriers to the absorption of drugs. *Daphnia* are a common model organism widely used in ecotoxicological testing^22,23^: they display high permeability and high sensitivity to compounds in their environment, which is an essential characteristic for drug screening^23-25^. Prior studies have demonstrated that the behavior of *Daphnia* is altered by chemicals, nanoparticles, pesticides, or bacteria products^22^, which allows for the dose-dependent tests. In contrast, *C. elegans* and *Drosophila* are non-aquatic species and therefore it is difficult to accurately profile dose-dependence of pharmacological perturbations due to high individual variation in effective drug consumption. Thus, the short-lived, fresh-water crustacean *Daphnia* offer a number of advantages over other common models of aging for screening of novel pharmacological agents.

Here we introduce a unifying framework for evaluating the effectiveness of anti-aging interventions using *Daphnia* as a model organism and a machine learning algorithm to build a predictive model with longitudinally tracked phenotypes. Specifically, we designed a scalable culture platform for longitudinal monitoring of *Daphnia*. With this platform, we tracked animals longitudinally in a cohort of *Daphnia* until their natural deaths by developing a computer vision algorithm that quantitatively extracts a representation of the location and behavioral parameters of individual animals in the culture tanks. We therefore used extracted features to train a supervised machine learning algorithm to predict their phenotypic age and compare them to chronological age. We found that our predictive model was able to accurately estimate phenotypic age that might reflect animals’ health state. We then evaluated the robustness of our model in such experimental conditions as drug or chemical treatment and examined how much these perturbations affect the animals’ healthspan. The developed analysis pipeline allows for quick and efficient tests for potential pharmacological candidates that increase the healthspan. The high-throughput, scalable, automated approach presented here enables the extraction of new behavioral features and training the model to evaluate the effect of perturbations on their behavioral outputs according to their experimental purposes.

## Results

### A scalable culture and high-throughput longitudinal phenotyping platform

Most current *Daphnia* behavior tracking platforms have been designed for short-term experiments (e.g., a few minutes to hours) with a small number of animals (usually 5∼10 animals per tank) for toxicological tests^22,26,27^. However, understanding the relationship between phenotypic changes and the aging process requires longitudinal monitoring with a large number of individuals due to the stochastic nature of aging processes. Neonates removal is an essential aspect of the long-term *Daphnia* culture. Since an adult female produces eggs every 3∼4 days until the death and neonates grow rapidly^28^, we should adequately separate neonates from mothers to maintain age-synchronized cohorts. However, this is still performed manually and is a labor-intensive and time-consuming task. To improve upon existing imaging systems for the long-term and a large number of animals, we engineered an integrated platform that provides 1) a long-term culture with large sample size, 2) scalability provided by making an individual culture tank a module, and 3) controllability of stimuli for profiling behavioral responses of daphnids by implementing the mesh-based tank design and an integrated behavioral assay platform (Fig. 1).

**Figure 1.**
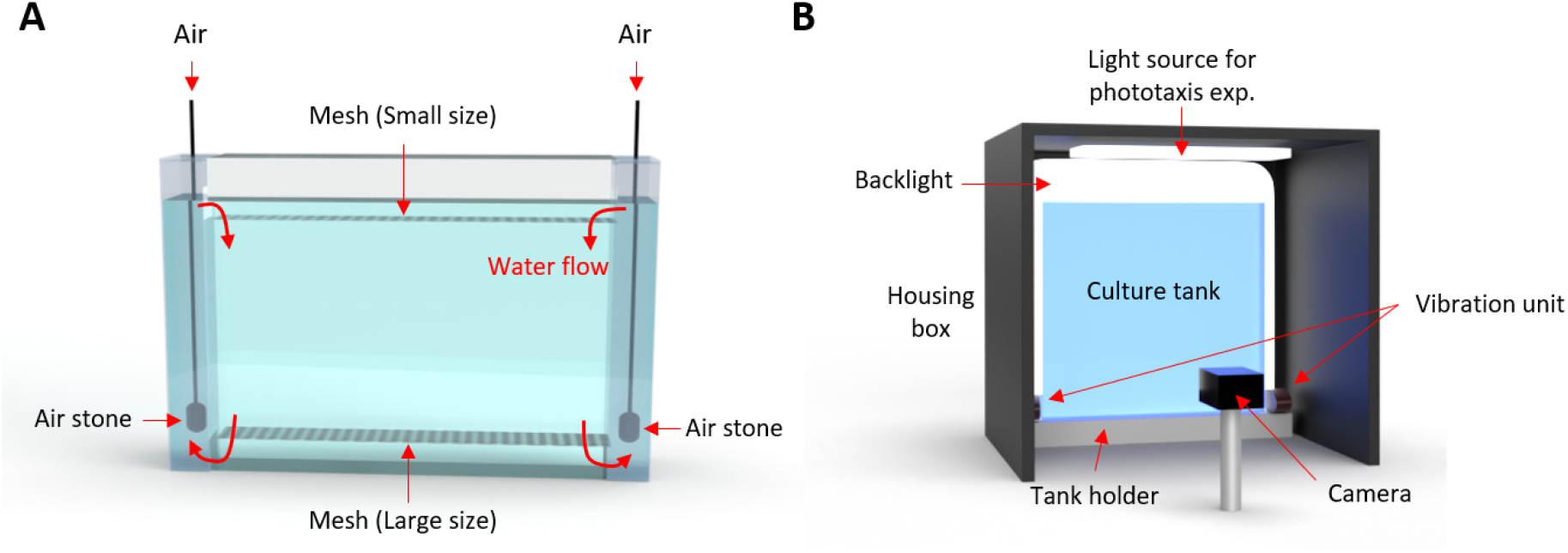
Our platform enables long-term culture and monitoring of daphnid behaviors. A-B) Schematic illustrating the customized culture and imaging setup. **A**) An individual tank. Air stones, connected to the air source, are located in two side columns to create an aerobic environment for daphnids and to separate neonates from mothers via two different sized meshes. **B**) Schematic of imaging setup. A tank is set into the imaging setup and recorded via a frontal camera in a computer-controlled environment. An even backlight illumination is used constantly. A housing ceiling light is used for the stimulated phototaxis. The scaffolding ensures an invariant tank placement.

To easily separate neonates from mothers and to monitor animals’ phenotypes, we designed a tank which consists of three modules; a housing tank, an insert, and a cap part with two different sizes of meshes (850 μ*m* mesh at the bottom of the insert and 300 μ*m* mesh at the bottom of the cap) (Supplementary Fig. 1A and C). To provide sufficient air to the daphnids because they thrive in an aeration condition, we created a continuous air-lift water flow through the two side columns which are separated from the insert tank where animals are housed (Supplementary Figs. 1A and B). Due to the effect of the continuous flow system, neonates are naturally passing through the large mesh at the bottom of the insert tank and get caught at the small mesh of the cap. Therefore, to separate neonates, we just need to remove and wash the cap. When we tested this platform, all 34 neonates were separated from the main tank within 5.5 min (Supplementary Movie 1). The geometry of the tank is optimized for the 1L volume of media to culture a large number of animals (The size of a housing tank: 23 cm (w) x 20.5 cm (h) x 4 cm (d)). *Daphnids* can swim freely in a relatively large arena (The size of an insert: 16.5 cm (w) x 14.5 cm (h) x 2.5 cm (d)). In addition, we can maintain and scale-up multiple different experimental conditions in a single experimental period (e.g., test different drugs, multiple drug concentrations or food level at each tank depending on the experimental design and progress) due to the modularity of the system. It also represents truly independent replicates.

Cognitive decline is one of the noticeable age-related changes observed across many species^29^. Since light and vibration are well-known stimuli to induce behavioral responses in *Daphnia*^*30*^, it would be essential to accurately control the light and/or vibration stimulus and monitor its responses with age to measure the effect of the perturbation on the cognitive ability. To allow precise temporal and strength control of stimuli and recording under the controlled environment, we developed an automated imaging setup (Fig. 1C). All parts (e.g., camera, lighting, vibrational motor and data organization) are integrated and controlled via Arduino board and a custom MATLAB GUI, which enables automatic experimental setup, controlling stimulus intensity and timing, and monitoring the phenotypic changes of the *Daphnia* as they age (Supplementary Figs. 2 and 3).

### A computer vision algorithm for extracting quantitative phenotypes

To extract multiple behavioral features quantitatively, there is a need for a robust analysis pipeline. Even though there is commercially available software for *Daphnia* behavioral monitoring, it has been applied to a limited number of animals in a tank (∼5 animals per tank) for a short period of time (a few minute to hours)^26^. To process the data from our high animal-density experimental conditions, we developed a custom MATLAB script that extracts various behavioral and morphological features to track phenotypic changes during the aging process. Briefly, we performed background subtraction and image segmentation to identify and determine the location of alive individuals (Fig. 2A, Supplementary Fig. 4 and Supplementary Note 2). Using a consensus approach informed by positional and morphological parameters, we separated the individual animals from other objects, such as dust or gunk, and performed tracking followed by measurements of the behavioral features (Fig. 2B).

**Figure 2.**
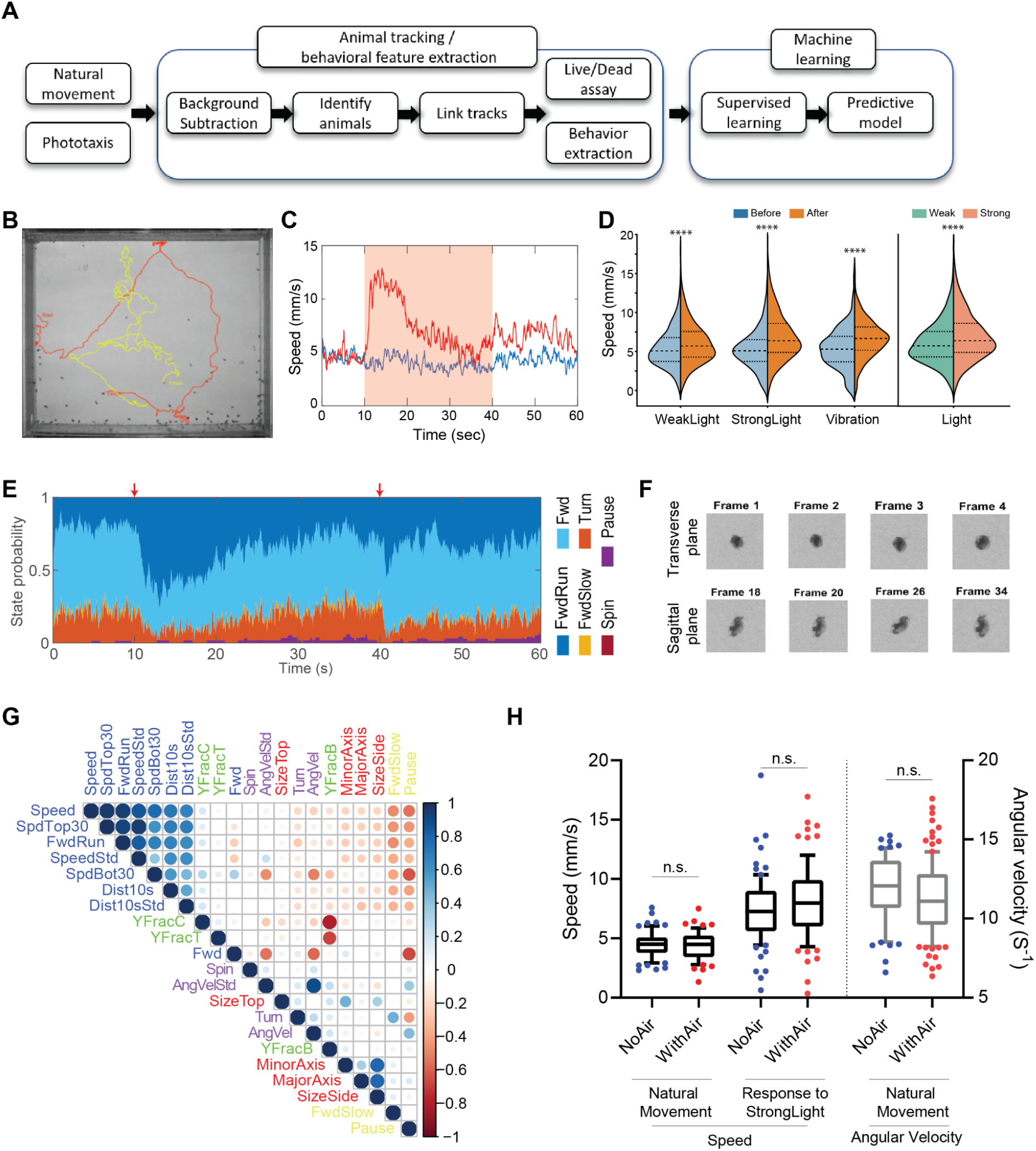
Quantitative behavioral analysis. **A**) A workflow of the behavior tracking and predictive model building. **B**) Two sample animal trajectories in a phototaxis experiment. **C**) Examples of extracted speed in (i) normal and (ii) 30s light-stimulus conditions. The shaded light red box indicates the timing of a light stimulus. D) Speed changes in response to various stimuli; weak light, strong light and vibration (Day 16). **E**) Descriptive behavioral features in 30s light-stimulus conditions (See Supplementary Table 1 for details). **F**) Example images of the animal for specific size extraction. Using circularity as a criterion, we distinguished the size of the transverse and sagittal planes (See methods for details). **G**) The correlation matrix of features, extracted from the natural swimming conditions (Using all longitudinally recorded videos). **H**) Comparison of two quantitative features, speed and angular velocity, between no air and continuous airflow conditions with the same animals at the same age. Statistical analysis: two-tail t-test (*<0.05, **<0.01, ***<0.001, ****<0.0001).

When we considered which features can be extracted, we started with parameters that had previously been associated with toxicological testing^22^. For example, the speed has been shown as one of the sensitive parameters in many *Daphnia* toxicology tests^22^. The vertical movement in a phototaxis experimental condition can be another useful feature. Therefore, we monitored animals’ movement in two regimes, natural swimming and stimulus-induced response, to examine age-related phenotypic changes in both cases. For example, the average swimming speed shows different patterns between these regimes. In the natural swimming mode (e.g., no external stimulus), we observe relatively consistent swimming speed. Yet, when animals are exposed to light, they show an obvious response: the average speed is increased dramatically as animals move up towards the light when the light is on (Fig. 2C). Eventually, the speed stabilizes at the level we observed prior to switching the light on. On the other hand, once the light is off, animals move towards the bottom of the tank (Fig. 2C, Supplementary Fig. 5, and Movie 2). Furthermore, our platform allows to control the brightness of light (Strong light: 2.65 ± 0.02 kLux and Weak light: 0.75 ± 0.01 kLux). Therefore, we tested how daphnids respond to different strengths of light stimuli. Lastly, since the vibration is known to evoke daphnids responses^30^, we monitored the animals’ responses to controlled vibrational stimulus. Figure 2D shows the distribution of speed before and after each stimulus. *Daphnia* clearly responds to each stimulus and shows graded responses to the different strengths of light. It indicates that the controlled stimuli in our platform can induce daphnids behavioral responses.

Several studies in various model organisms have shown that morphological parameters such as muscle mass, body weight and size are associated with lifespan or diseases^31-33^. Thus, we also extracted morphological features related to animals’ size (Fig. 2D and Supplementary Movie 2). Furthermore, we created a list of locomotor classifications such as ‘forward fast running (FwdRun)’, ‘forward swimming (Fwd)’, ‘Forward slow swimming (FwdSlow)’, ‘turning (Turn)’, ‘spinning (Spin)’, and ‘pause’ which involve basic locomotor actions of *Daphnia*. The analysis pipeline computed the per-frame probability of each locomotor description. Figure 2E shows that most of the time, young and healthy animals show forward swimming (either FwdRun or Fwd). The next-most common behavior was turning, in which *Daphnia* made a large, rapid change in orientation. But when the light stimulus was delivered, most animals changed their behavioral status from either forward swimming or turning to forward fast running. It indicates that our descriptors of behavior are robust in reflecting animals’ behavioral changes. In total, we extracted a set of 21 quantitative features focused on natural swimming behaviors and 12 additional features related to phototactic response (**Supplementary Table 1**). To understand the relationship among extracted features, we calculated the correlation matrix (Fig. 2E). As expected, features in different categories are largely independent of each other, but ones in the same category (e.g., the forward movement features, turning features, pause mode, morphology-related features, and vertical location-related features) are more correlated.

Since we created a continuous flow for 24 h to create air-driven circulation in the tank, except when we recorded the video, we tested whether this continuous flow in itself alters the animals’ behavior. There were no statistical differences between no-air and with-air conditions (Fig. 2F) in both speed and angular velocity. It suggests that our continuous flow is nonrestrictive enough not to alter daphnids’ swimming patterns. Furthermore, to evaluate the performance of the automated algorithm, we segmented the 1 min video into four overlapping segments (e.g., 30 s each: 0∼30s, 10∼40s, 20∼50s, and 30∼60s, respectively) and compared the results of processing of each fragment for the features including speed, and angular velocity. We found that the replicated data are not significantly different amongst the fragments (Supplementary Figure 6). It indicates that our algorithm was able to measure physiological features in a robust, reproducible manner.

### Highly controlled survival probability across replicates in the platform

The platform allows to count the number of alive animals for survival assay to build a lifespan curve and evaluate the effect of perturbations on lifespan. However, the manual live/dead assay is a heavily labor-intensive task and there is no available algorithm for *Daphnia* lifespan assay based on our knowledge. Therefore, an automated counter is required to build lifespan curves from our large-scale high-throughput longitudinal experiments. However, since animals often overlap or touch each other in the high-density population case, it is not easy to accurately segment and detect all individuals to count them. Since our algorithm could not count all live animals perfectly (e.g., count 75.62 % of animals with 0.29 % error (mislabeling)), we developed a hybrid approach. A MATLAB GUI is used to manually curate the counts (Supplementary Figure 7). Since more than 75% of animals are already automatically and correctly counted, this pipeline makes counting much faster than entirely manual and much more accurate than entirely automated.

Reproducibility in the new platform is one of the critical factors in finding robust age-related biomarkers and then conducting phenotypic screens for pharmacological interventions that promote healthy aging. We tested whether three independent cohorts could show reproducible lifespan curves. Figure 3A shows that reproducibility between cohort experiments is high (e.g., no statistical difference with the log-rank test; median lifespan: cohort 1 = day 53, cohort 2 = day 52, cohort 3 = day 57). Not surprisingly, individual trial reproducibility is relatively low (Supplementary Figure 8). It serves to inform us that the general health of the natural variants is well controlled in our platform if the size of the sample is large enough (> 90 animals per cohort). It indicates the feasibility of the platform to test age-related interventions.

**Figure 3.**
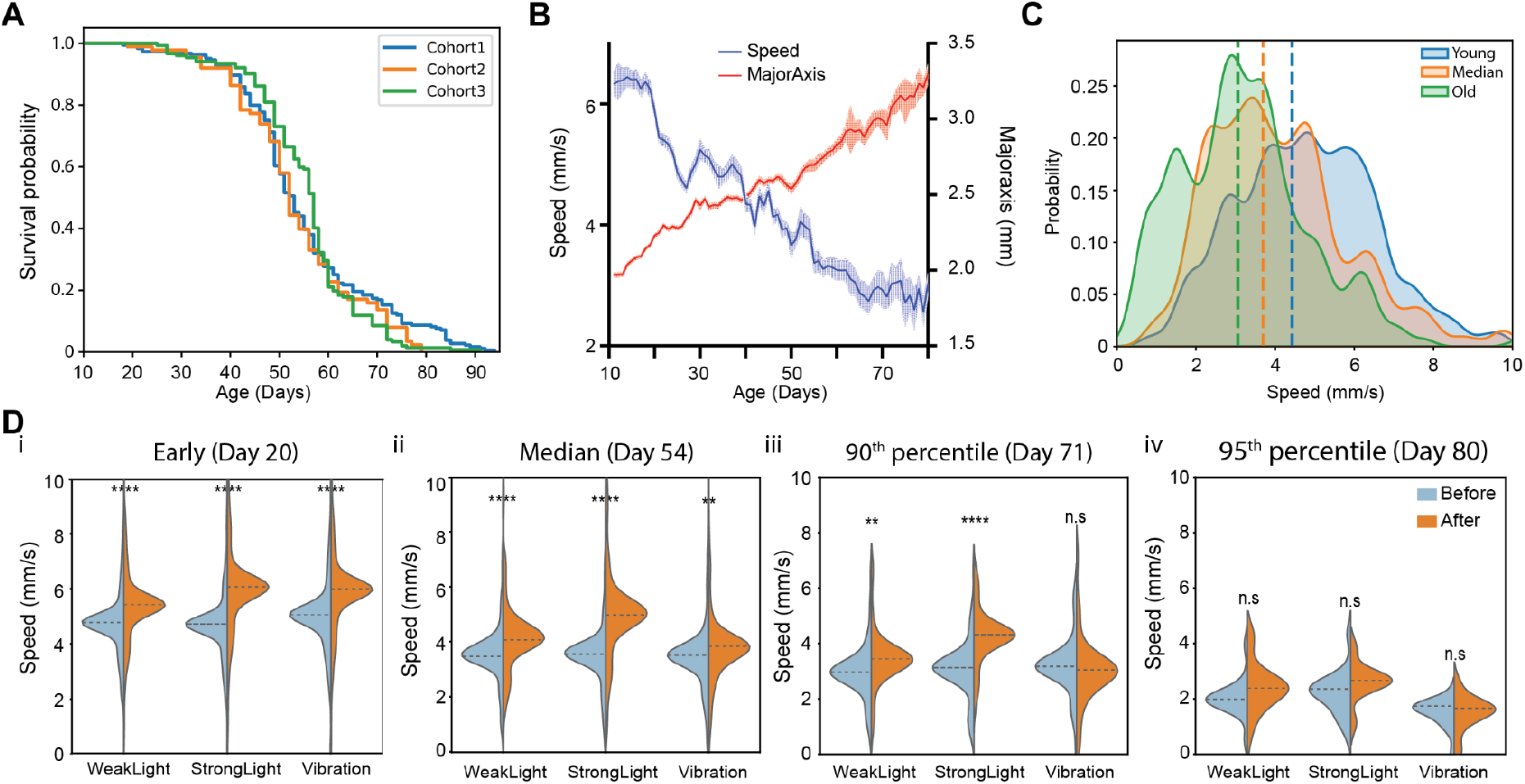
Age-related phenotypic changes. A) Lifespan curves for three control experiments (Cohort 1: n = 184, Cohort 2: n = 88, Cohort 3: n =152). B) A lifetime population average of speed (blue) and body size (red) of *daphnids* in a control experiment. Error bars are SEM. **C**) Density plot of speed at three ages: young (blue - 20th percentile), median (red - 54 day old) and mature (green - 80th percentile of the lifespan), **D**) Response to stimuli (weak light, strong light, and vibration) speed before (blue) vs after stimulus (orange) at four ages (i: young, ii: median, iii: mature, iv: old). Two-tailed t-test (** p-value < 0.01, **** p-value <0.0001)

### Age-related phenotypic changes

Age in animals is accompanied by a decline in locomotion and decreased responses to various stimuli, which are direct measures of healthspan^34^. Motility and responses to controlled stimuli were thus assessed over the entire adult lifespan of animals. As expected, the average swimming speed decreased with age, while body size increased at the population level (Fig. 3B). Behavioral features display variation at the individual level, but the population-based average value shows the trend of age-related changes. It is known that *Daphnia*, like most other crustaceans, continues to grow their entire life^28^. As we mentioned, both light and mechanical stimuli can induce *Daphnia* responses. Thus, we set on to quantify this effect using the speed parameter. Figure 3D shows the change of speed before and after the three different stimuli over the lifetime (we selected four timepoints; young age (Day 20), the median of lifespan curve (Day54), 90^th^ percentile of the lifespan curve (Day 70) and very old age (Day 80)). When animals are young (Figure 3D i and ii), they significantly respond to all three types of stimuli, even though strong light induces faster movement than weak light. When animals get older, the distributions of speed for both before and after stimuli are gradually decreased. Specifically, at age day 71, animals still clearly respond to both light stimuli, but vibrational stimulation does not induce significant responses (Figure 3D iii). At old age (Day 80), animals do not show significant responses (Figure 3D iv). This establishes that our platform is sufficiently sensitive to capture the change of animals’ behaviors with age.

### The effect of metformin treatment on both the lifespan and phenotypic features

We next asked how pharmacological perturbations influence lifespan as well as behavioral decline during the aging process. While several potential anti-aging drugs have been suggested in various model organisms, they have not been well tested in *Daphnia*. As a proof of concept of the suitability of the developed model for the measurement of phenotypic age in perturbed conditions, we used one of the well-known anti-aging drugs, metformin, in *Daphnia magna*. Metformin is a drug commonly prescribed to treat patients with type 2 diabetes and several studies suggest that metformin can increase the lifespan of various model organisms^11,12,35^.

We first determined the long-term effects of four doses of metformin in female daphnids (1 mM, 1 μ*M*, 0.1 μ*M* and 0.01 μ*M*). The age when drug treatment is started may also be an important variable to consider. Since, in mice, early life metformin treatment can extend mean lifespan while late-life treatment failed to increase lifespan^36^, we started to apply metformin when animals were transferred to our culture platform after they were fully developed (Day 12∼13 when they start to produce progenies). Unsurprisingly, it is observed that a significant decrease in lifespan for daphnids cultured at a high dosage of metformin compared to those at the lower concentrations or control condition (all animals were dead within 2 days in the toxically high 1mM metformin concentration), following known trends in other model organisms^11^ (Fig. 4A). 1 μ*M* metformin is also toxic and significantly shortens the median lifespan of female daphnids by 14.5 % (χ^2^ = 20.42 and p-value < 0.0001 in log-rank test). However, the significantly shifted lifespan at the low concentrations of metformin is not observed (Control vs. 0.1 μ*M*: χ^2^ = 1.078 and p-value = 0.297 / Control vs. 0.01 μ*M*: χ^2^ = 2.092 and p-value = 0.148 in log-rank test).

**Figure 4.**
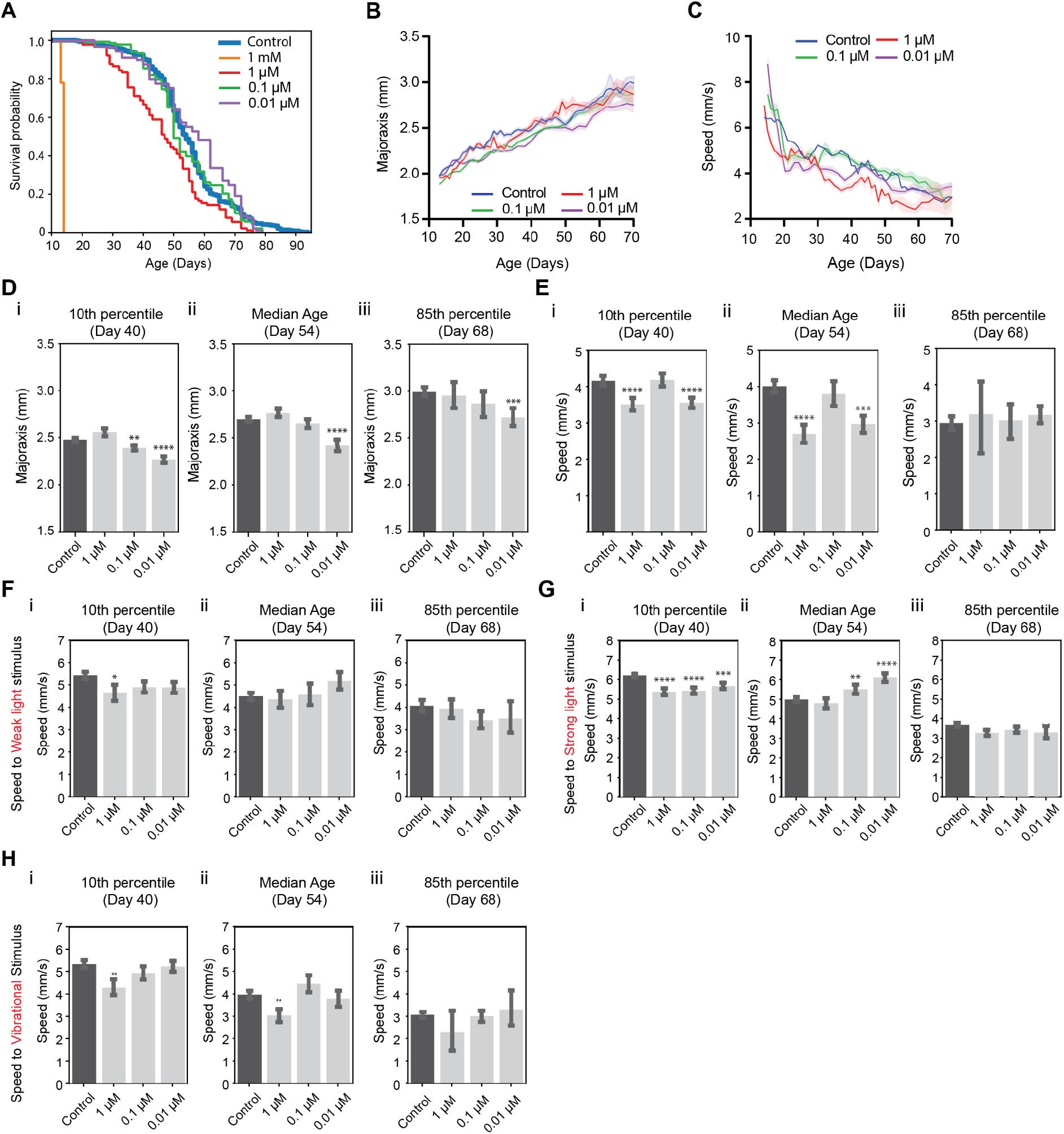
Behavioral changes and longevity in metformin-treated animals. **A**) Survival curves (Control: n = 424, Met 1 mM: n =90, Met 0.1 μM: n = 138, Met 0.01 μM: n = 89), **B**) average body size and **C**) average speed in the natural condition (SEM error bars), D) Body size and **E**) Speed at different ages. F, G, H) Response to stimuli (weak and strong light, vibration) at different ages. All tests compared to control via two-tailed t-test, *<0.05, **<0.01, ***<0.001, ****<0.0001.

To investigate whether behaviors and morphological features are impacted by metformin treatment, we first plot the changes in body size and speed by age with the result of the control group (Figs. 4B and C). Interestingly, the body size of 0.01 μ*M* metformin treatment animals is smaller than the control one over almost an entire lifespan (Fig. 4D). In mice, metformin-treated male mice were lighter than control animals^11^. In 1 μ*M* and 0.01 μ*M* the movement in the normal condition is slower than in control animals until the median age (Fig. 4E). Based on lifespan assay, 1 μ*M* dose is toxic, likely causing the slowdown. 1 μ*M* metformin-treated animals also show reduced responses to external stimuli (Fig.4F-H). On the other hand, the reduced speed of 0.01 μ*M*-treated animals might be caused by a small body size. In a steady environment, smaller animals move slower, but when responding to stimuli they might be able to react more actively due to the effect of the drug (Figs. 4F-H).

### Quantitative behavioral metrics can estimate physiological ages using machine learning

As individual features might be correlated with chronological age, we sought to determine the degree to which individual features could correlate with chronological age. We performed a simple linear regression on each feature for the measured age and evaluated its performance. The major axis parameter shows the best correlation with age, followed by sagittal plane body size (Supplementary Table 2). While several single parameters were somewhat correlated with chronological age, we can expect that we could build a more accurate predictive model using combined extracted features as input.

With the ability to extract multiple quantitative behavioral features, we asked whether these could be compiled into a single predictive model to estimate the physiological ages of individuals using the integration of phenotypic parameters as input features and chronological age as output labels. We built models using five types of machine learning models – LASSO (least absolute shrinkage and selection operator), Elastic Net, Random Forest^37^, Gradient Boosting and SVM (Support Vector Machine) – and quantify the extent to which the model fits the data in both natural (e.g. no external stimulus) and stimuli-induced conditions (Supplementary Table 3). As expected, the multivariate models show better predictive performance than one using univariate, with lower error, higher r-squared value. Particularly, the random forest model had the best accuracy with the highest r-squared value and lower mean error than other models for both natural swimming and stimulus-induced experimental data (Figs. 5A, B and Supplementary Fig. 9). A decision tree-based Gradient boosting approach also shows very similar performance as Random Forest. This might be that tree-based ensemble models such as Random Forest and Gradient Boosting can represent complex interactions among features, which linear regressions, such as LASSO and Elastic Net, cannot do^37^. Unsurprisingly, the model built using stimulus-induced phenotypic features shows better predictive performance than one using natural swimming conditions. This can be attributed to the stimulus-induced behaviour being more reflective of age-related performance, and possibly to the fact that we use more features to represent the stimulus-induced swimming dataset (Fig. 5D, E and Supplementary Table 3).

**Figure 5.**
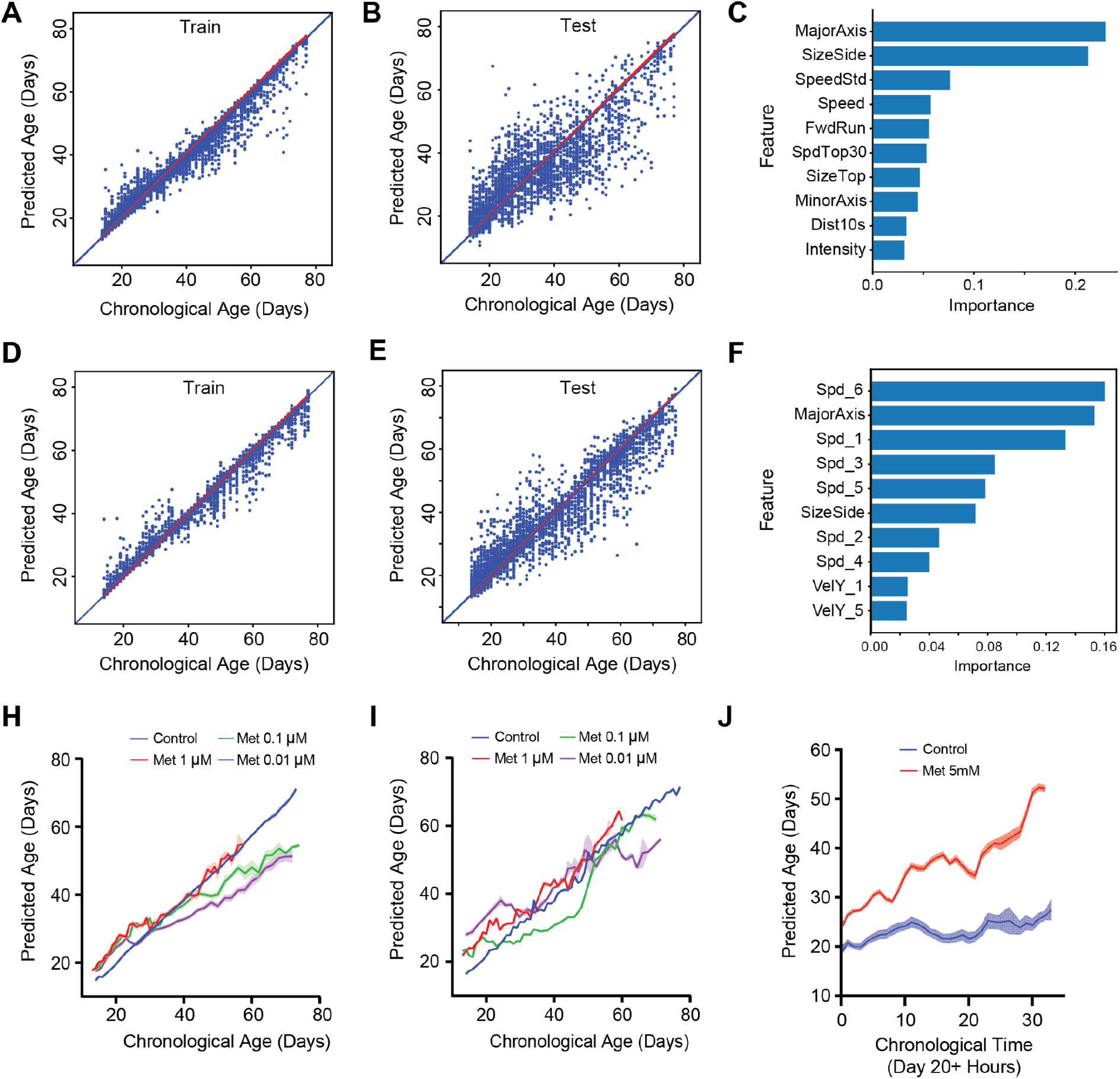
The predictive model using Random Forest. A-F) The model built by Random Forest predicts ages from A, D) training and B, E) testing data sets (data was randomly divided 7:3, separated by individual trajectories) of A-B) the natural (12k individual trajectories) and D-E) stimulus-induced condition. The diagonal blue line indicates a theoretically perfect prediction. (See Supplementary table 3 for the accuracy test). C, F) Importance of the top 10 features derived by Random Forest for C) the natural and F) stimulus-induced condition (See Supplementary table 3 for the definition of features). H, I) Comparing predicted age vs chronological age for control and metformin-treated animals using the model developed by the natural condition (H: natural and I: stimulus-induced condition). Error bars are SEM. J) Comparing predicted age vs. chronological age (Day 20) for control and 5mM metformin-treated animals in every hour recorded stimulus-induced phenotypes (control: n = 22 and 5 mM: n = 19).

One of the benefits of the random forest method, in addition to a highly predictive model, is that we can rank relevant features from Random Forest using the permutation feature importance analysis, i.e. by calculating the incremental error resulting from the feature being excluded from the model (Figs. 5C and F). As the result of a simple linear regression for the individual feature shown, size-related features were identified as the important features, followed by the speed feature. Interestingly, the lists of important features of the models are different based on the experimental conditions (natural vs. stimulus swimming). Specifically, features indicating the responsiveness to the stimulus (speed or y-directional velocity) come out as important in the stimulus swimming compared to natural swimming. It might imply that parameters showing cognitive ability are highly related to the health status of animals. features in the stimulus swimming model compared to the natural swimming one. It might imply that parameters showing cognitive ability is highly related to the health status of animals.

The predictive accuracy of our established model suggests that our model may be useful for evaluating the lifespan effects of multiple interventions in *Daphnids* many days before their death. Specifically, we hypothesized that the difference between predicted age and chronological age might be representative of phenotypic age, where healthier animals would be estimated to be younger than their chronological age. To test this idea, we calculated this difference in metformin-treated animals using the predictive models (e.g., using both models built by normal condition data and stimulus condition data) (Figs. 5H and I). Both models estimated the age of low (0.1 μ*M* and 0.01 μ*M*) metformin concentration-treated animals to be smaller than that of control animals. On the contrary, 1 μ*M* metformin-treated animals predicted ages are usually higher than the control one. If we calculated the slope of the simple linear regression (forced x- and y-intercepts are zero), the control group shows almost one (0.9639 ± 0.0005 by natural condition and 0.9833 ± 0.0007 by stimulus condition) (Supplementary Table 4). However, the low concentrations of metformin treatments show smaller slope values than the control one. On the other hand, a high concentration of metformin shows slope values larger than 1, which indicates that the rate of the aging process is differently estimated in our model. With the important features for the models, we can assume that lower behavioral activity in 1 μ*M* contributes to the old phenotypic ages by the both models (e.g., no significant size difference between Control and 1 μ*M* condition (Fig. 4D)). In the lower concentrations, both smaller body sizes and behavioral responses may contribute to the younger phenotypic age in both cases. It demonstrated that we can use the predictive model to estimate animals’ phenotypic

Furthermore, this model allows us to quantitatively evaluate the toxicity of drugs at an early time point. Figure 5J shows the phenotypic age of both control and 5 mM metformin-treated animals. In this experiment, we monitored animals’ phenotypes every hour in both groups in the stimulated motion conditions and then estimated the phenotypic age. All animals in the 5 mM treated group died within 1.5 days. Even a few hours later, the predicted age of 5 mM metformin-treated animals is higher than the control one. Then the difference of estimated age is getting larger. It opens up the possibility that our model may be used to test the toxicity of drugs on animals’ health within a few hours.

### Estimated physiological ages in various perturbed conditions

The key application of our predictive model is to quantify the health of individuals at an early age to test the effects of interventions which perturb animals’ lifespan and healthspan. This avoids the need for monitoring phenotypic changes over the entire lifespan to evaluate the efficacy of drugs. To validate this idea, we tested several chemicals and quantified the effects of chemicals using our model. Various studies have reported the effect of chemicals on *Daphnia* physiology, such as movement or heart rate in a short time period. Among them, ethanol was reported to reduce the heart rate of *Daphnia*^*38*^. When we treated daphnids with 1 ∼ 4 % ethanol, the behavior was dramatically changed within minutes (Figs. 6A-C). For example, within 5 min of 3% ethanol treatment, animals’ movement speed sharply decreased (Fig. 6C and Supplementary Video 4). When phenotypic age was calculated for these ethanol-treated animals, they appeared to be older than the control animals (e.g., control: 16.29 ± 0.89 days, 3% EtOH 5min: 18.05 ± 0.57 days, p-value < 0.001). On the other hand, caffeine was reported to increase the heart rate of *Daphnia*^38,39^. Interestingly, our model indicates that the 0.8mM caffeine-treated animals appear younger than the controls, which is statistically significant (p-value <0.01) (Fig. 6D). To further evaluate our model, we tested solutions of Ficoll, which makes higher viscosity of culture media. Since aquatic animals swim slowly in high viscous media, it is expected that animals’ behavior in Ficoll-mixed media is inhibited compared to standard ADaM culture. Naturally, our model estimated the much older phenotypic ages for these animals than the control animals (Fig. 6E) and reflected the progressive change with the concentration of Ficoll. Lastly, we tested fluoxetine which has been reported to make a slight change in phototactic responses on *Daphnia*^*26*^. However, in our analysis, even though the phenotypic age in the phototaxis case is estimated older than the control one, it did not show any significant changes in both natural and phototaxis conditions (Fig. 6F).

**Figure 6.**
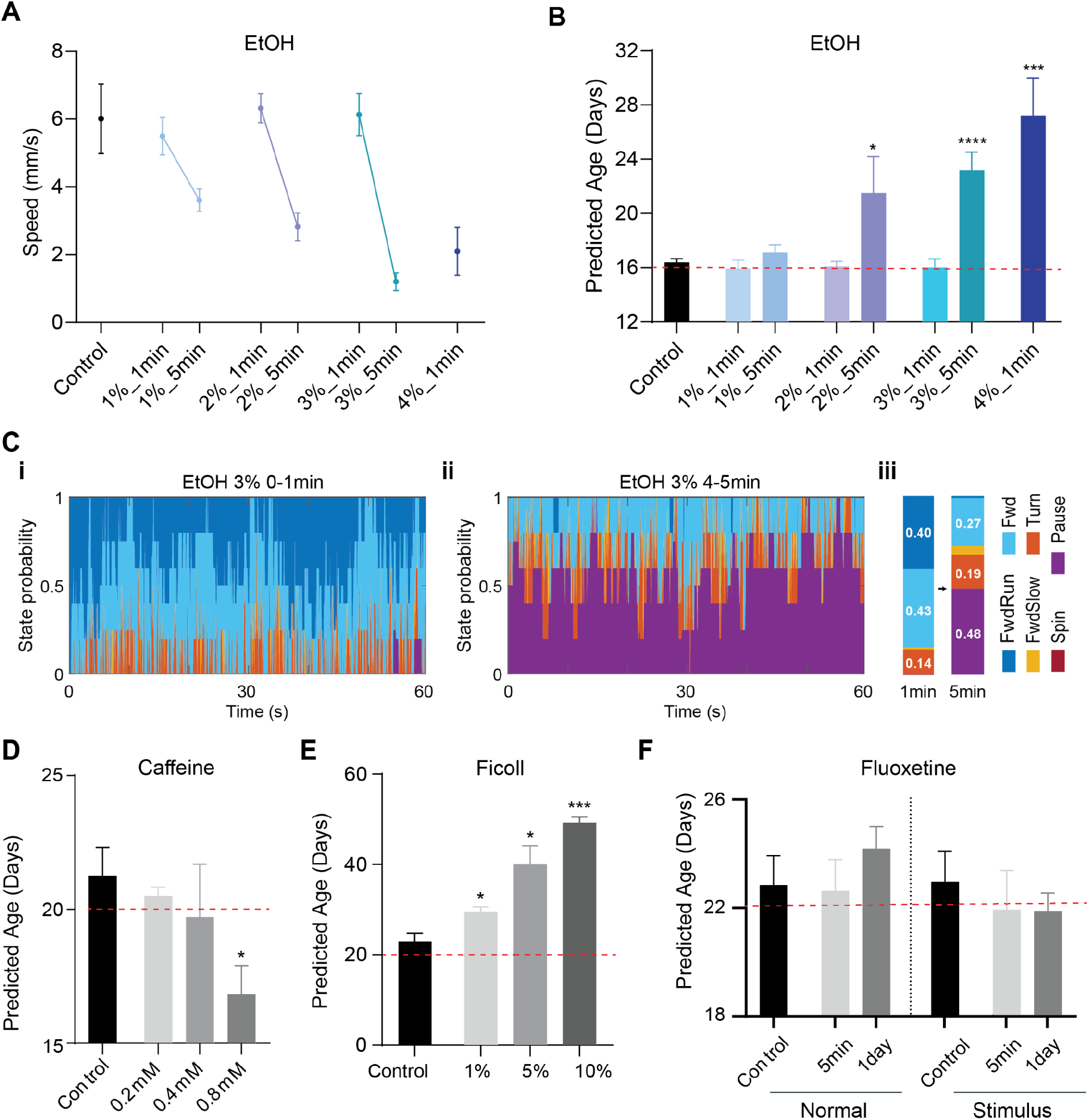
Estimated phenotypic ages in various interventions in young adult daphnids. The red dash line indicates the chronological age. Mann-Whitney test (p-value: * < 0.05; ** <0.01, and *** < 0.001). Error bars are SEM. A-C) The effect of ethanol treatment (n = 5 for each condition). **A**) Speed changes on ethanol, **B**) Predicted (Phonotypic) age. The chronological age of the tested animals is indicated by the red dash line. **D**) caffeine (n = 3 for each condition), **E**) Ficoll (n = 5 for each condition), and **F)** Fluoxetine (n = 11);

Furthermore, we used smaller sizes of the tank for the short-term chemical tests to examine the possibility of reducing the quantity of chemicals and drugs required and the overall cost of the screen. Specifically, we used the original size of the tank (1L volume) for fluoxetine, a medium size of the tank (400ml volume) for ethanol and Ficoll, and the small size of the tank (100ml volume) for caffeine (Supplementary Figure 9). In all cases, the estimated ages for control are very similar to their chronological ages (Figs. 6B and D-F). It indicates that our framework is not specific for a particular design of the experiment. It can be more widely applicable for various shapes of culture tanks depending on the experimental purposes. In conclusion, these test results indicate that our model can predict how old individual animals appear to be and it is sensitive enough to distinguish the effect of various interventions on the health of animals.

## Discussion

The quantitative phenotypic biomarker for the aging process allows for measuring the health status of animals and it would be essential to develop new anti-aging pharmacological strategies. The discovery of such pharmacological interventions in aging requires high-throughput screening strategies. However, the majority of screens performed in model organisms so far are rather low-throughput only with live/dead assays. Given the complexity of the aging process, the aging biomarkers would be multifaceted, not a single feature. Therefore, measuring various features longitudinally in a high-throughput manner is a key to establish the meaningful metrics of biomarkers for the aging process and successfully transfer pharmacological approaches to the clinic. For *Daphnia*, although several systems for monitoring behaviors exist, most of them are dedicated to measuring 1-2 features in a short period of time. In this study, we demonstrated an integrated platform that allows for the longitudinal culture and monitor of *Daphnia* phenotypic changes with minimum experimenters’ efforts in a high-throughput manner. The platform’s inherent modularity can accommodate various experimental conditions at the same time. With our customized software, we can extract a variety of behavioral and morphological features systematically. Thus, our automated platform and analysis pipeline provides high-content phenotypic measurements of *Daphnia*.

Many biological processes relevant to the aging process can be accounted for as phenotype-based changes. Using our platform, we monitored age-related phenotypic changes and lifespan in control and metformin-treated animals until their natural deaths. We found that *Daphnia* shows the shift of lifespan curves depending on metformin dosage, suggesting that *Daphnia* is sensitive to examine dose-dependent drug tests. In other words, the whole-animal screens provide information about the important toxicological, pharmacokinetic, and pharmacodynamic activity of the screened compounds at an early stage. Thus, it is possibly reducing attrition rates for the downstream phases of the drug development process. We also observed that various behavioral features declined during the aging process, as like other model organisms^11,40^. This observation points that our platform and pipeline can establish the areas in which *Daphnia* can complement existing models through improvements in sensitivity, cost, or efficiency for the aging research and phenotypic screening for the anti-aging drug discovery.

We first built a model as the phenotypic clock in *Daphnia* from an ensemble of quantitatively extracted features. We demonstrated the feasibility of this clock by examining phenotypic changes across different chemical perturbations on a short-term scale (a few minutes to 1 day). Then, the use of our predictive model can quickly evaluate the effect of these chemicals on the aging process. It suggested that the integration of phenotypic features enables the construction of an estimator for phenotypic ages to explore the relationship between perturbations and the aging process. However, the performance of the predictive model can be more improved. First of all, we can improve the accuracy of the tracking algorithm using convolutional neural networks instead of the conventional image process technique that we used here. Due to the overlapping of animals in a high-density population, our current algorithm could not track all animals properly during entire recorded frames. A convolutional neural network may provide a better solution for object segmentation and tracking problem. Second, the predictive accuracy could be improved by adding more sample numbers, especially at the older ages, because we have a smaller number of trajectories in old ages compared to the younger ages. Lastly, the model could get an advantage from the incorporation of additional input features. Specifically, by measuring performance on additional cognitive tasks in our platform, such as chemosensation in *Daphnia*^*30*^, we would deeply understand the functional correlates of cognitive changes with normal aging and predict future cognitive impairment.

The major limitation of phenotypic-based screens is that the mechanism of action of drug hits is unknown and there is a possibility to have false positives due to compounds that target genetic mechanisms such as protein synthesis, which affect the assayed phenotype but are not specific enough to be used as drugs. This limitation would overcome in combination with innovative approaches to genome, transcriptome, proteome, fluorescent markers, or molecular reporters. By adding multifaceted information, we can investigate integrative biology at high resolution across multiple organ systems, cellular, and genome levels at multiple time points during the aging process. Specifically, the transparent body of *Daphnia* makes real-time observation of its cell biology and physiology straightforward. With this observation, we can understand how particular organs or systems fail with age. For example, we observed the decreased swimming ability with age that might indicate the degradation of muscle and the decreased performance in phototaxis with age that might indicate the defeats in cognitive ability. By measuring muscle loss and detecting damage in neuron number, synaptic integrity, and neurotransmitter, we will find the causality factor for the age-related phenotypic decline. Thus, use those high-dimensional multifaceted data to construct dynamic networks that will provide a better understanding of what extent individuals age differently and enable assessment of potential interventions by providing more information-rich readout.

Using a short-lived model organism, an automated video monitoring platform and phenotypic profiling pipeline, and machine learning algorithms, we show the possibility of that the non-invasive phenotypic measures could be used in perturbational studies to understand whether a perturbed condition is effective to delay aging at an earlier than its death. In theory, our approach could be adapted to predict phenotypic age for other model organisms. The ability to predict the health status of animals enables us to conduct rapid screens for anti-aging drugs. We envision that our framework can greatly expand the repertoire of not only high-throughput behavioral measurements but also deep pharmacological profiling in *Daphnia*. As future work, it is capable of high-throughput screening large libraries of small molecules for their effects on particular traits of interest, such as extension of healthspan and/or lifespan. Specifically, we can re-examine drugs from the “DrugAge” database, which compiles results on lifespan effects from > 500 distinct compounds in more than 20 species^41^. Since very few drugs are tested across multiple species, having most of these compounds profiled in the same organisms during the aging process will create the benchmark for the development of pharmacological strategies that extend the period of healthy life and eventually prevent or reduce the onset of age-related phenotypic changes or diseases.

## Methods

### 1. Population culture

*Daphnia magna* animals were used in all assays. IL-MI-8 heat-tolerant clone was obtained from the Ebert lab at the University of Basel, Switzerland stock collection originating from a pond in Jerusalem, Israel. To collect synchronized cohorts, neonates born within 1 ∼ 2 days were separated from the mothers and their sex was determined at day 8 ∼ 10. To minimize the effect of maternal age, we used a large range of maternal age from Day 11 to Day 90. All mothers are also well-fed and cultured at the same conditions (25°C incubator). We used females for all experiments in this study to avoid sex-dependent differences. Collected females were cultured until when animals start to lay first-neonates (approximately day 12 ∼ 13 at 25°C) and then 40 ∼ 50 animals were randomly assigned to one of our developed culture tanks. All cultures were maintained in ADaM water^42^ at the 25°C incubator, exposed to a light cycle of 16 light hours followed by 8 dark hours, and fed every other day the suspension of the green alga, *Scenedesmus obliquus*, at a concentration of 1 × 10^5^ cells/ml (for 1 animal/10ml density, amount of food prorated by population). Every fourth day the water was changed and offspring were removed manually until animals transferred to the culture platform. The air flow in the tank was regulated by a pressure gauge. The operating range of pressure was 1 ∼ 1.5 psi depending on the number of connected tanks.

### 2. Imaging tank fabrication

A custom-designed transparent tank for *Daphnia* was made from acrylic sheets (McMaster-Carr, USA). A single tank setup consists of three pieces; a housing tank, an insert, and a cap (Supplementary Figure 1). Based on the dimension of the tanks, acrylic sheets were cut using a laser cutter. The cut pieces were bonded by acrylic cement (SCIGRIP 16, USA). The housing tank is partitioned into three parts as follows; two side columns were used for air generation and the middle part for the insert used for recording. The bottom of the insert is made from a 850 μ*m* mesh to separate progeny from their mothers. The bottom of the cap is made from a 300 μ*m* mesh to prevent the progeny from getting into the insert.

### 3. Video acquisition and stimuli control

Three videos were taken once a day. The recording was made at a rate of 25 fps using a 1.3 Megapixel monochrome CMOS camera (DCC3240M, Thorlabs) coupled with an optical lens (MVL8M23, Thorlabs). For the longitudinal tracking data, the video was recorded every day until all animals died. A white LED background light (LightPad 930, ArtoGraph) was provided to create an even illumination into the entire tank and a white LED strip was used for light stimulus. Using the neutral density filter, we minimize the intensity of the backlight (0.16 ± 0.01 kLux). To record the video, we moved the tank to the imaging setup, turned off airflow to create static conditions. The animals were then left to acclimate for 2 ∼ 3 min with the backlight before the actual recording began. We filmed two videos; 1) 1 min video to capture natural swimming behavior and 2) 2 min video to monitor behavioral responses to controlled stimuli. All videos were stored using unique file names identifying the cohort, recording time and the experimental condition. The following two recording regimes were used: 1) natural swimming with no extra light stimulus; and 2) stimuli-induced swimming (e.g., each stimulus was delivered for 10s and use 20s as the interstimulus interval). The intensity of strong light stimulus at the top of the tank was 2.65 ± 0.02 kLux and one at the bottom of the tank was 0.32 ± 0.01 kLux. The intensity of weak light stimulus at the top of the tank was 0.75 ± 0.01 kLux and one at the bottom of the tank was 0.10 ± 0.01 kLux. The strong light stimulus is approximately three times brighter than the weak light stimulus. The tank was first covered by a dark housing box (Figure 1C) to control the amount of stimulus light accurately by minimizing the effect of ambient light and also induce animals to assemble at the bottom of the tank as the baseline before the light stimulus.

### 4. Animal detection and tracking

We performed background subtraction and image segmentation to determine the location and identity of animals. Specifically, the background was calculated as the average of every 25^th^ frame (25 fps) and subtracted from each frame to remove all stationary objects (e.g., dead corpses and dust). We then segmented moving objects to determine the potential boundary of the animals and the identities of the animals were determined by a consensus informed by positional and morphological parameters. All segmented animals in each frame were parameterized by the centroid position, size, major- and minor-axis length. The two body-size features of *Daphnia* (e.g., body size of transverse vs. sagittal plane) were determined by the circularity of segmented *Daphnia*:

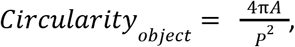

where A is the area and P is the perimeter. Here the size of the transverse plane was classified when the normalized circularity value is larger than 0.92 and the size of the sagittal plane was classified when the normalized circularity value is smaller than 0.58. These thresholds were empirically selected.

The code is available in GitHub.

(Platform control GUI: https://github.com/dydals320/ImagingGUI

Behavior tracking and analysis code: https://github.com/dydals320/DaphniaBehAnalysis)

### 5. Predictive model

Longitudinal tracking data is intrinsically imbalanced due to the death of animals. To overcome this limitation for the regression performance, we used a minority oversampling technique, SMOGN (Synthetic Minority Over-Sampling Technique for Regression with Gaussian Noise)^43^. We tested several machine learning models such as LASSO, Elastic Net, Random Forest, Gradient Boost, and SVM in the scikit-learn module of python^44^. The performance of the models was determined with the adjusted R^2^ and RMSE values. There were 12,356 total data points (individual trajectories) for the normal behavioral condition and 16,517 data points for the stimulus condition in the control group. Missing data were replaced by the mean value for a given feature for a given age group.

## Acknowledgments

The authors would like to thank Camille Homa for the help of *Daphnia* husbandry, Doosan Jung for the discussion of ML model and Wil Ratzan for critical reading of the manuscript. This work was supported by the Paul G. Allen Frontiers Group and McKenzie Family Charitable Trust.

