## Supplemental Methods for ""Smart Tanks": An Intelligent High-Throughput Intervention Testing Platform in *Daphnia*"

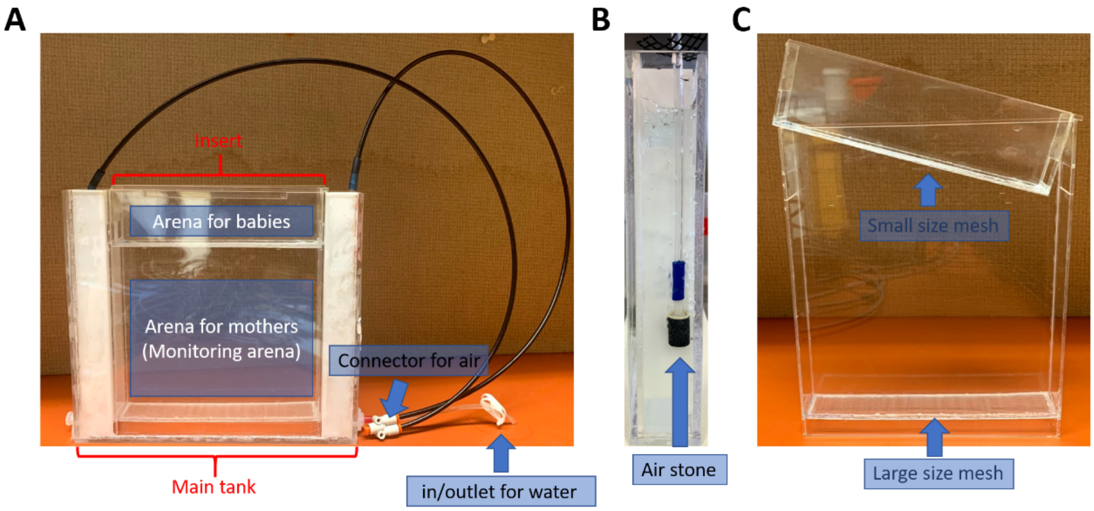


**Supplementary Figure 1.** Images of the tank module.

Picture of an actual tank, consisting of three parts: 1) (A) Main tank has (B) two side columns in order to create air-bubble, 2) (C) the insert has large mesh at the bottom to separate progenies from mothers, and 3) the cap which allows for capturing separated progeny using a fine mesh.

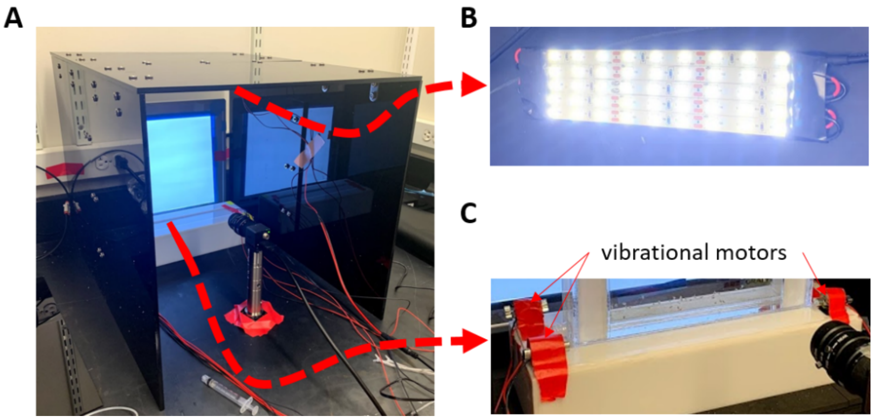


**Supplementary Figure 2.** Imaging setup

A) The backlight panel placed behind the tank, the camera positioned at the front of the tank, B) the white LED light strip is on top of the tank (underneath the black housing box) for phototaxis assay. C) The slot bracket allows for the placing of a tank at the exact same position for each imaging session and four vibrational motors are attached at the corner of the tank holder to generate controlled vibrational stimuli.

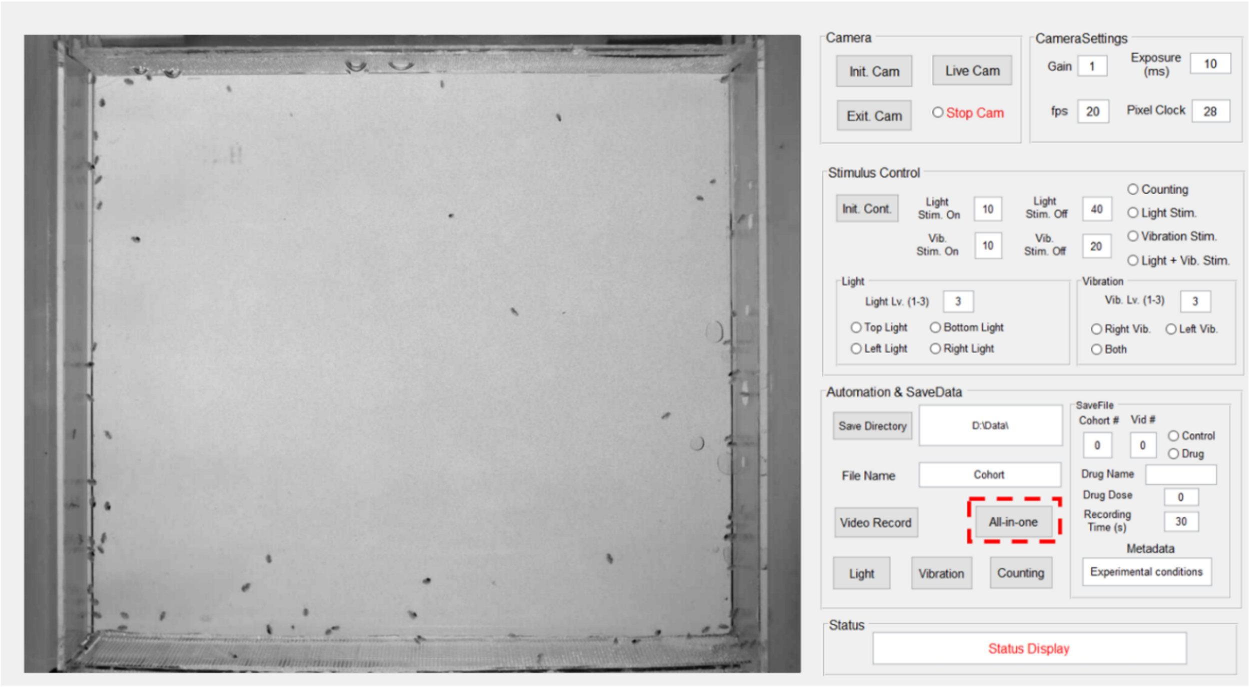


**Supplementary Figure 3.** The interface of the user GUI

The left window shows the currently monitored field of view. Right panels allow experimenters to manually control camera and stimulus (duration and intensity of both light and mechanical stimuli), save files with typed metadata, and view/take records of a tank in real-time. All these steps can be scripted and automatically performed by clicking an 'All-in-one' button. The current GUI status is shown at the bottom of the status panel.

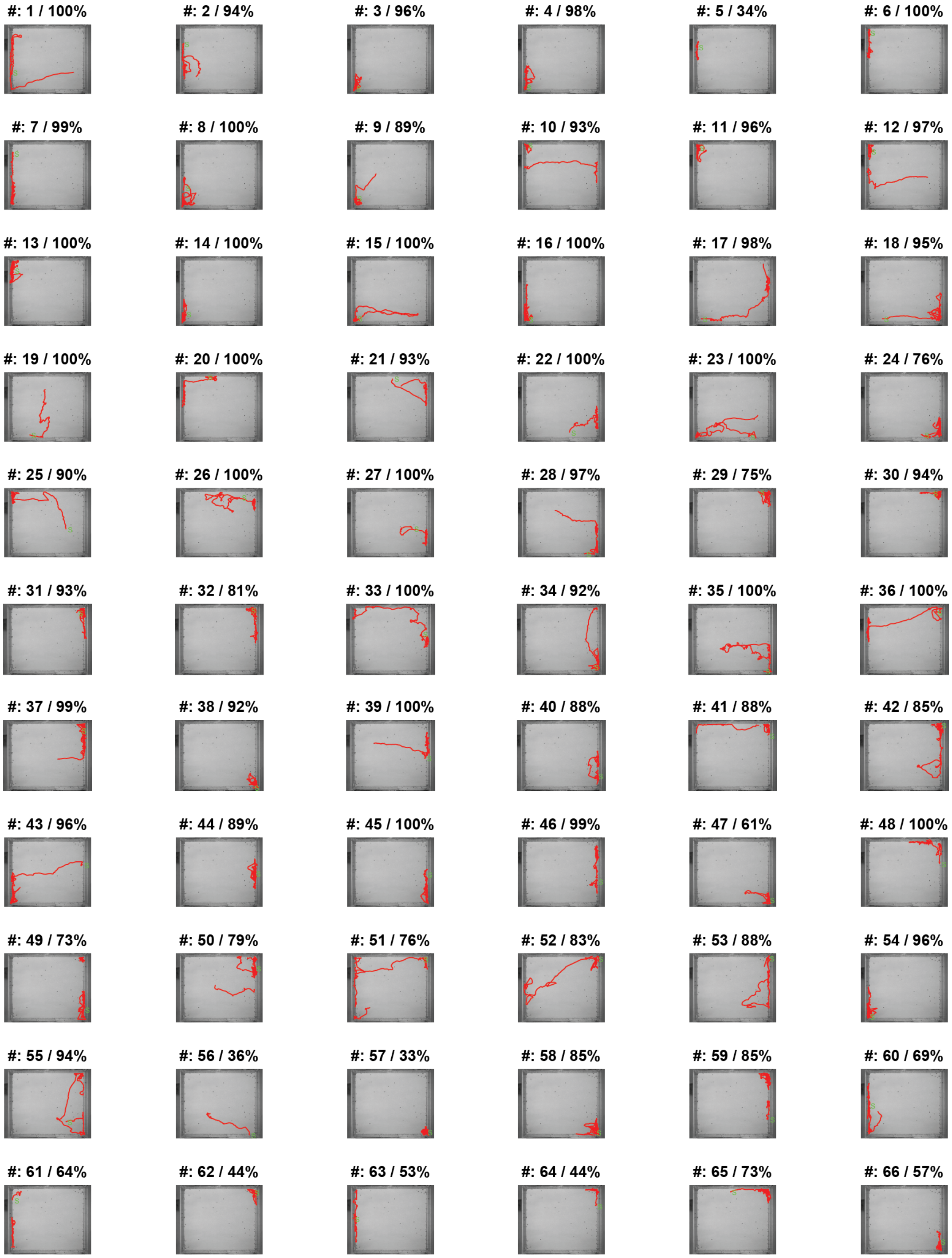


**Supplementary Figure 4**. Example trajectories from a video

Individual trajectories from Day 14 control 1 min long video (78 live animals). The red line indicates the automatically tracked trajectory of the animal. The green letter 'S' indicates the starting point. The percentage is showing how many frames are tracked. We extract behavioral parameters from each trajectory that got tracked during more than 20 s (33 %) in the 1 min video.

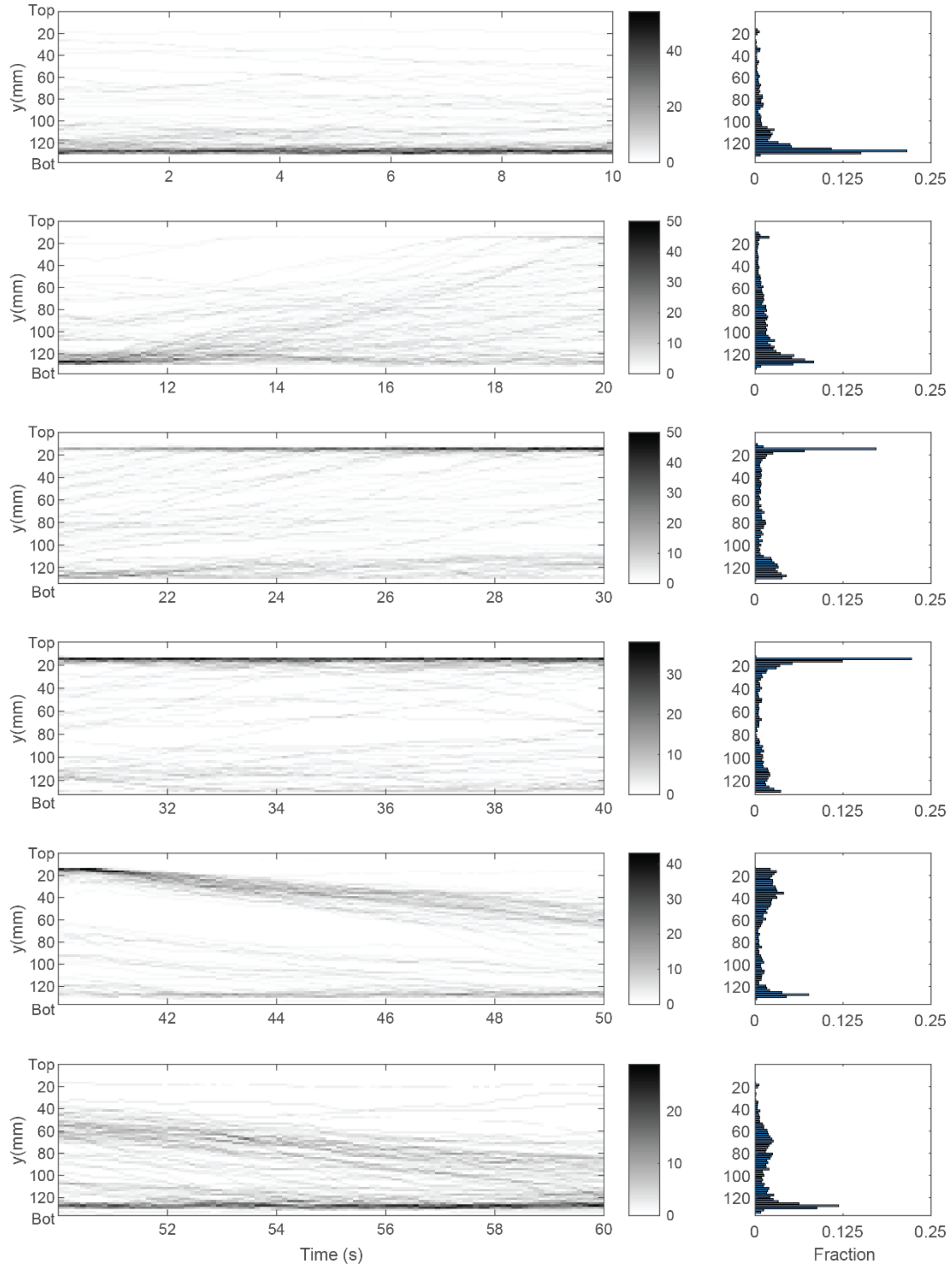


**Supplementary Figure 5.** Vertical movement of daphnids in the phototaxis experiment.

During this 1 min experiment, the light gets turned on at 10 s and then off at 40 s. Plots on the left show the vertical density of animals in the tank through time. Plots on the right show the histogram of animals' location. At the beginning, most animals stay at the bottom of the tank. When the light is on (at 10 s), most animals start moving up towards it and remain at the top of the tank. Once the light is off (at 40 s), animals start returning to the bottom of the tank.

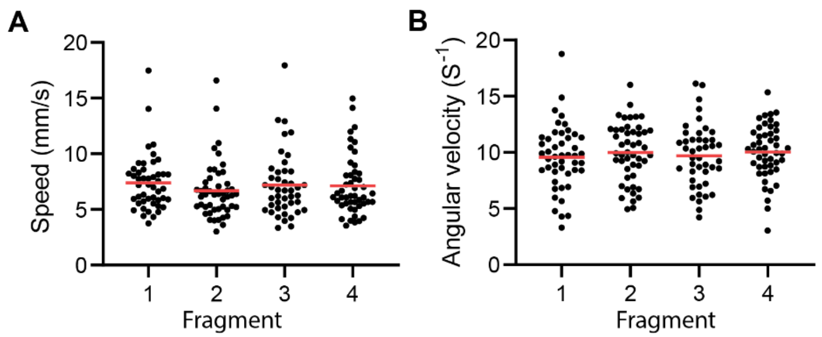


**Supplementary Figure 6.** The evidence of the measurement consistency

Comparison of speed and angular velocity across the 30 s long fragments of 1 min long video. All repeated data are not significantly different from one another. ANOVA test.

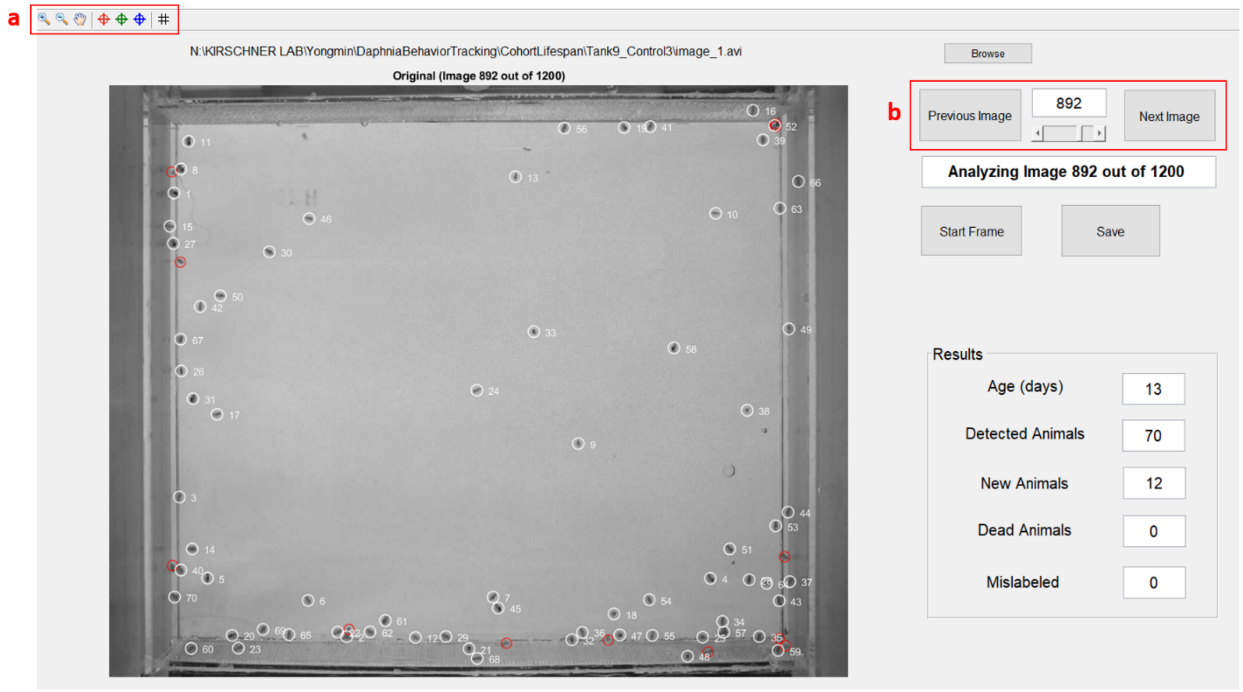


**Supplementary Figure 7.** MATLAB GUI for animal counting

After loading a video, the algorithm found a frame that might have the most identified animals and show the number of detected animals (labeled as a white circle and the ID number in the image) in the result panel. Using options in section a (red box), users can zoom-in / -out the image and add undetected animals (labeled as a red circle in the image), dead, or mislabeled animals. To change the frame, users can use the options in section b (red box); moving a frame by clicking the previous image or next image button or jumping ten frames by using a slide.

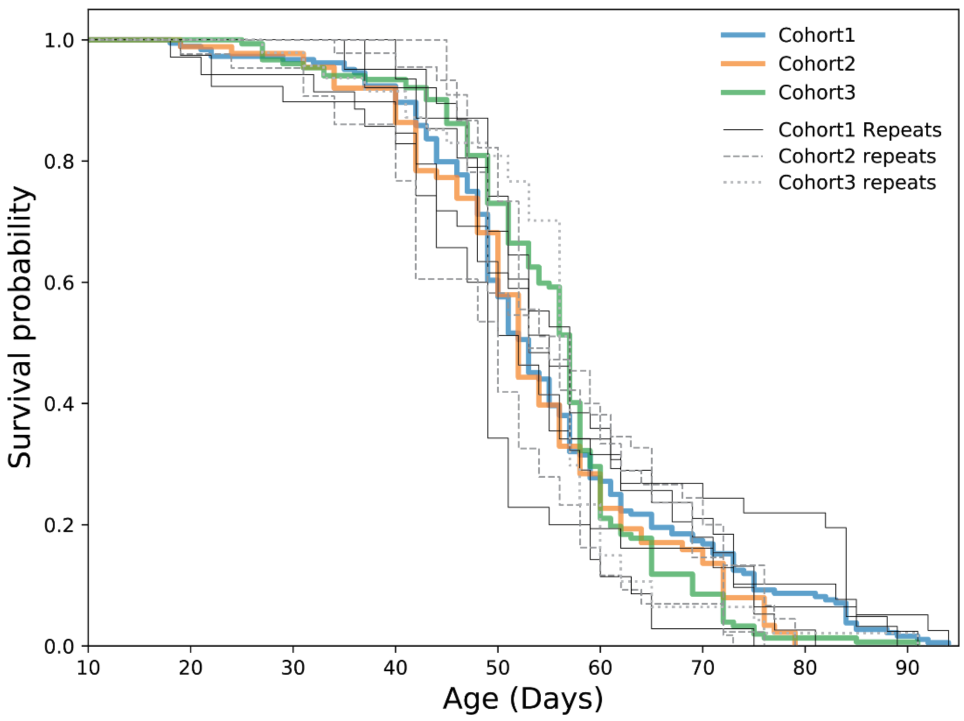


**Supplementary Figure 8.** All survival curves (control condition).

The survival curves set displays the total range of observed lifespan for each of the 10 experimental replicates (think black lines) from three cohorts at the control condition. Thick color lines show the averaged longevity of each cohort. Overall, the average longevity across the cohort experiment did not differ (log-rank test).

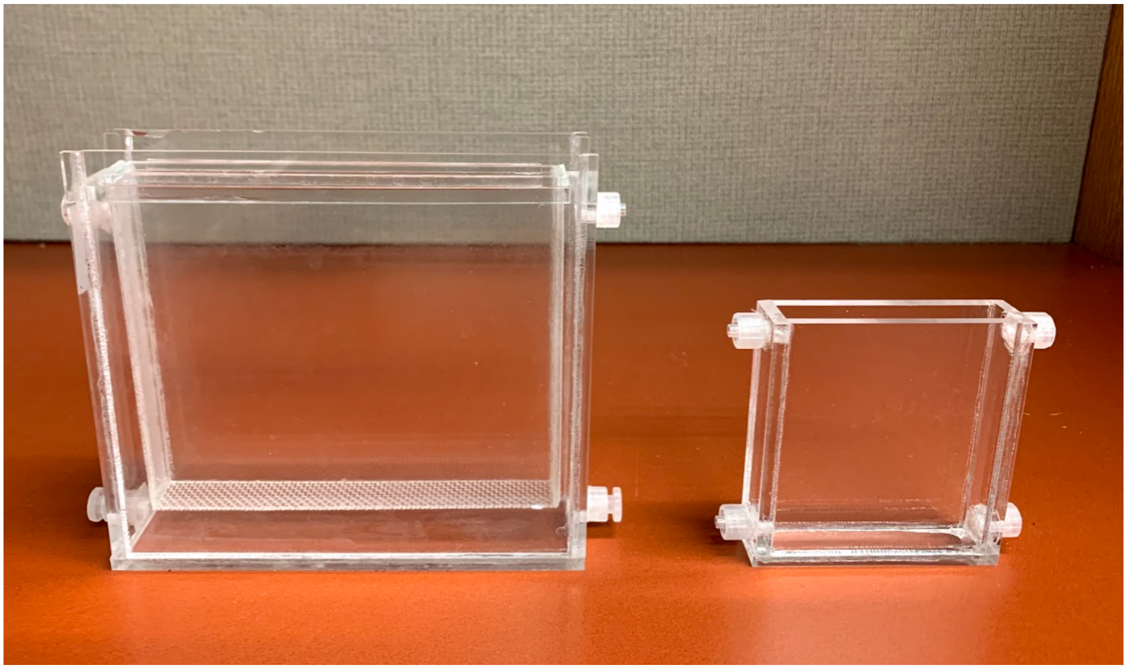

**Supplementary Figure 9.** Smaller tanks for short-term drug tests

Picture of the tanks for short-term drug tests. The volume of the left one is 400 ml (15 cm (w) x 13.5 cm (h) x 3.5cm (d)) and the right one is 100 ml (7.5 cm (w) x 8.5 cm (h) x 2 cm (d)).

**Supplementary Table 1**. A list and description of quantitatively extracted features

<table>
<tr><th>Experimental conditions</th><th>Category</th><th>Feature</th><th>Description</th><th># of features</th></tr>
<tr><td rowspan="10">Both natural- and stimulus-swimming conditions</td><td rowspan="3">Swimming ability</td><td>Speed (SpdTop30, SpdBot30)</td><td>SpdTop30 (or SpdBot30) is consist of the fast<br>(or slow) 30th percentile speed.<br>+ standard deviation of speed</td><td>4</td></tr>
<tr><td>Swimming distance for 10s</td><td>+ Standard deviation</td><td>2</td></tr>
<tr><td>Angular velocity</td><td>+ Standard deviation</td><td>2</td></tr>
<tr><td rowspan="4">Morphology</td><td>Size of transverse plane</td><td></td><td>1</td></tr>
<tr><td>Size of sagittal plane</td><td></td><td>1</td></tr>
<tr><td>Major axis</td><td rowspan="2">Major / minor axis of sagittal plane</td><td>1</td></tr>
<tr><td>Minor axis</td><td>1</td></tr>
<tr><td>Location in a tank</td><td>YFraction_Bot/Cen/Top</td><td>How long Daphnids stay on each location.<br>Equally divide a tank as 3 section vertically</td><td>3</td></tr>
<tr><td>Descriptive features</td><td>forward fast running (FwdRun), forward swimming (Fwd), 'Forward slow swimming (FwdSlow), turning (Turn), spinning (Spin), and pause</td><td>FwdRun: 50% faster than total average speed (8.25 mm/s)<br>FwdSkiwL 50% slower than total average speed (2.75 mm/s)<br>Turn: Trajectory's curvature is larger than 0.25<br>Spin: keep turn for 5 s</td><td>6</td></tr>
<tr><td colspan="4"></td></tr>
<tr><td rowspan="2">Only for stimulus condition</td><td rowspan="2">Swimming ability</td><td>Spd_1, 2, 3, 4, 5 and 6</td><td>Calculate speed using 10s-time window for each stimulus<br>1: before weak light stimulus<br>2: after weak light stimulus<br>3: before strong light stimulus<br>4: after strong ligth stimulus<br>5: before vibrational stimulus<br>6: after vibrational stimulus</td><td>6</td></tr>
<tr><td>VelY_1, 2, 3, 4, 5 and 6</td><td>Calculate y-directional velocity (Vertical movement) using 10s-time window for each stimulus<br>(condition is same as Spd)</td><td>6</td></tr>
</table>

**Supplementary Table 2.** Results of using a simple linear regression to calculate adjusted R-squared, p-value, and mean squared error (RMSE) for individual features as a function of chronological age based on the data on natural swimming in control animals. We performed a linear regression on each features for age with a training dataset and evaluated its accuracy on a test dataset (data were randomly divided 7:3 as train and test set).

| **Feature** | **Adjusted** $R^2$ | **RMSE** |
|---|---|---|
| Major axis | 0.42 | 11.47 |
| Body size (Sagittal plan) | 0.36 | 11.97 |
| SD of Speed | 0.22 | 13.26 |
| Speed | 0.17 | 13.69 |
| SpeedTop30 | 0.20 | 13.39 |
| SD of distance | 0.16 | 13.70 |
| FwdRun | 0.16 | 13.75 |
| Minor axis | 0.16 | 13.75 |
| Distance for 10s | 0.12 | 14.04 |
| Pause | 0.06 | 14.48 |
| FwdSlow | 0.06 | 14.50 |
| SpeedBot30 | 0.05 | 14.63 |
| SD of Angular velocity | 0.04 | 14.69 |
| Body size (Transverse plan) | 0.03 | 14.71 |
| Angular velocity | 0.02 | 14.84 |
| YFractionCenter | 0.01 | 14.91 |
| Intensity | 0.01 | 14.93 |
| Turn | 0.00 | 14.96 |
| YFractionBottom | 0.00 | 14.94 |
| YFractionTop | 0.00 | 14.98 |
| Fwd | 0.00 | 14.96 |
| Spin | 0.00 | 15.07 |

**Supplementary Table 3**. Model prediction accuracy on test and testing data set

1. Training Data set

| | Natural Swimming (No stimulus) | | Phototactic responses (Light stimulus) | |
|---|---|---|---|---|
| **Model** | Adjusted $R^2$ | RMSE | Adjusted $R^2$ | RMSE |
| LASSO | 0.571 | 9.627 | 0.617 | 10.709 |
| Elastic Net | 0.577 | 9.566 | 0.616 | 10.709 |
| Random Forest | 0.634 | 6.369 | 0.866 | 6.371 |
| Gradient Boost | **0.865** | **2.018** | **0.942** | **3.821** |
| SVM | 0.570 | 9.662 | 0.613 | 10.858 |

2. Test Data set

| | Natural Swimming (No stimulus) | | Phototactic responses (Light stimulus) | |
|---|---|---|---|---|
| **Model** | Adjusted $R^2$ | RMSE | Adjusted $R^2$ | RMSE |
| LASSO | 0.547 | 9.669 | 0.634 | 10.472 |
| Elastic Net | 0.547 | 9.940 | 0.634 | 10.472 |
| Random Forest | 0.654 | 8.200 | 0.838 | 6.982 |
| Gradient Boost | **0.672** | **8.429** | **0.892** | **5.730** |
| SVM | 0.538 | 10.083 | 0.631 | 10.568 |

**Supplementary Table 4.** Calculated slope from the simple linear regression (forced x- and y-intercepts are zero).

| Condition | Natural | Stimulus |
|---|---|---|
| Control | $0.9639 \pm 0.0005$ | $0.9833 \pm 0.0007$ |
| Metformin 1 $\mu M$ | $1.0410 \pm 0.0049$ | $1.1600 \pm 0.0036$ |
| Metformin 0.1 $\mu M$ | $0.9280 \pm 0.0042$ | $0.9264 \pm 0.0030$ |
| Metformin 0.01 $\mu M$ | $0.8228 \pm 0.0040$ | $0.9450 \pm 0.0042$ |

**Supplementary Note 1 – Platform design documentation**

The following documentation is a overview on how the entire platform was built and operated for the set of experiments shown in the article. By design, our system is modular and scalable. As a result, it can be customized depending on the user's requirements.

**1.1. Tank design**

Dimensions for the tank (A: Housing tank, B: Cap and C: Insert). All dimensions are given in millimeters. Sections through the device in front, bottom and side views are shown.

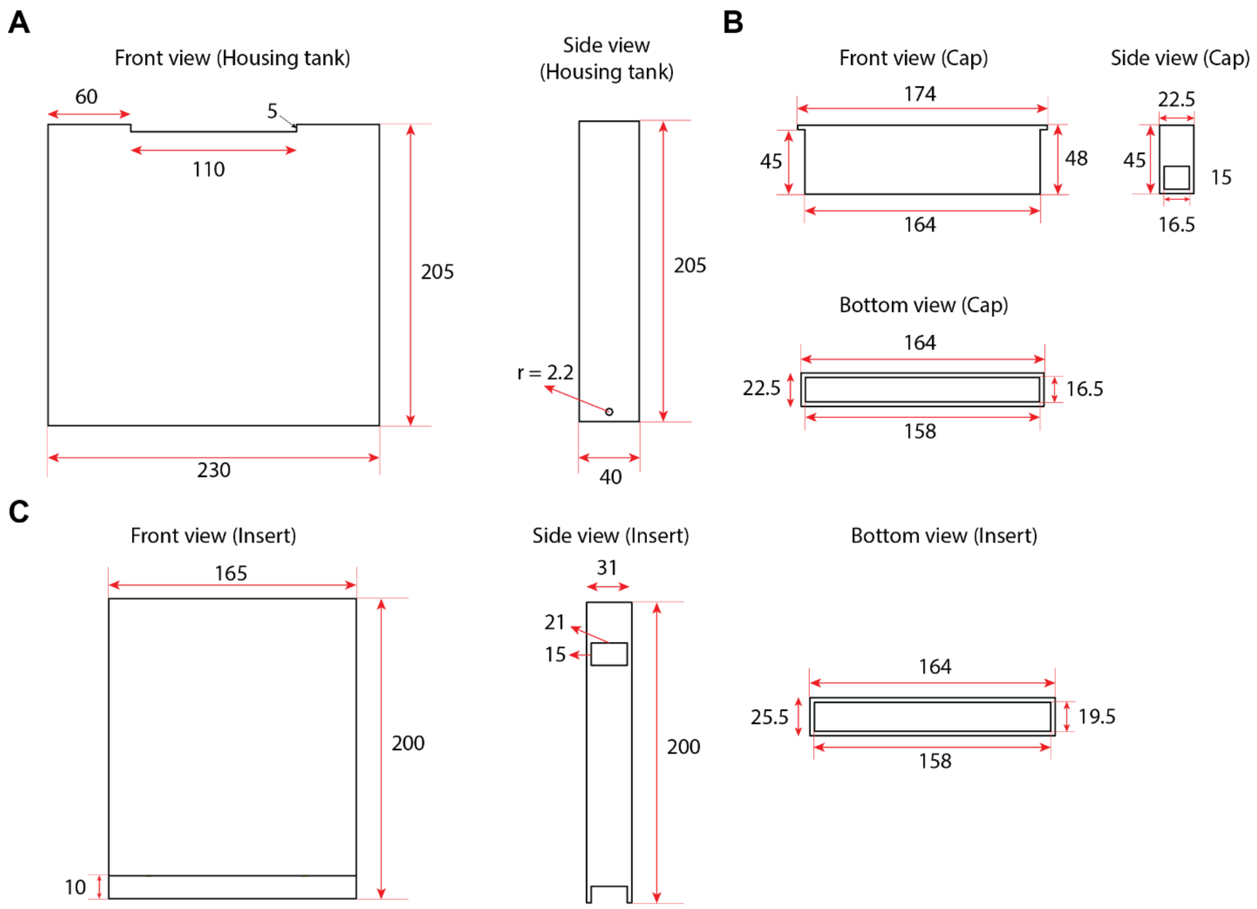

#### 1.2. Platform part list

| Category | Subcategory | Item name / Description | Vendor | Part number | Quantity | Note |
|---|---|---|---|---|---|---|
| **Tank system** | Tank | Clear Cast Acrylic Sheet, 12" x 24" x 3/16" | McMaster-Carr | 8560K219 | Depending on the number of tank | For housing tank |
| | | Clear Cast Acrylic Sheet, 12" x 24" x 1/16" | McMaster-Carr | 8560K172 | | For insert and cap |
| | | Cast Acrylic Sheet, 12" x 24" x 1/8", Color: White | McMaster-Carr | 8505K742 | | For the partition |
| | | SCIGRIP 16 Acrylic Cement | Amazon | B005ZH31W2 | 1 | |
| | | Female Luer Thread Style Cap | FTLLP-6005 | Nordson Medical | Pack of 100 | For outlet of the tank |
| | | Male Luer Integral Lock | XMTLL-6005 | Nordson Medical | Pack of 100 | |
| | Air control | Air Stones | Amazon | B076S3D75C | Pack of 12 | |
| | | Air Filter | McMaster-Carr | 8395K11 | 1 | |
| | | Air tubing | McMaster-Carr | 5548K82 | 100ft | |
| | | Compact Compressed Air Regulator | McMaster-Carr | 9892K11 | 1 | |
| **Imaging system** | Imaging | Camera / High-Sensitivity USB 3.0 CMOS Camera | Thorlabs | DCC3240C | 1 | |
| | | 8 mm EFL, f/1.4, for 2/3" C-Mount Format Cameras | Thorlabs | MVL8M23 | 1 | |
| | | Rosco Roscolux 398 Neutral Grey | stagelightingstore | Rosco Roscolux Sheet R398 | 1 | ND filter |
| | | Artograph LightPad, | Amazon | B01EX2LWTI | 1 | 12 x 12 inch |
| | Control & Automation | Arduino uno | Arduino | ARDUINO UNOREV3 | 1 | |
| | | LED Light Strip | Amazon | B075RYSHQQ | 1 | |
| | | Vibration Motor | Amazon | B07RGX3R1Q | Pack of 5 | Vibrational stimulus |
| | | Acoustic Damper | Amazon | B01M1EQHDO | 1 | Put the underneath of tank holder |

**Supplementary Note 2 – Phenotype tracking algorithm**

### 2.1. Overview

Daphnia phenotype tracking was performed using a multistep process beginning with raw video data. All analyses were performed in MATLAB 2020a using custom-written scripts, based in part on the previous algorithms[1] to segment video frames and identify continuous centroid paths of individual daphnids. Each frame was turned into a binary image using adaptive background correction and thresholding method. The resulting binary image was analyzed using a consensus approach informed by positional and size parameters to separate the target objects (adult Daphnia) from other objects and then performed tracking followed by measurements of the morphological and behavioral parameters.

### 2.2. The workflow of the algorithm

1) Setup parameters depending on the experimental conditions in the *DaphniaPhenotyping* (Main function).
2) Background subtraction and object segmentation and detection in the *DaphniaTracker.*
*3)* Extract various phenotypic parameters in the *DaphniaSegmentTracks.* Users can tune the feature extraction setting parameter in the *InitialSettings*.
4) Visualize the extracted features via *Ethogram* and *DaphniaDensity*.

**Supplementary Video 1.** Video shows how neonates (n = 34, day 1 - 3) are separated through mesh and continuous flow in the tank. Separated neonates are captured at the cap (top small size mesh). 10x playback.

**Supplementary Video 2.** Video shows how the algorithm tracks the centroid and detects the body size of the animal. Real-time playback.

**Supplementary Videos 3.** Animals' responses to 30s of light stimulus between 10th and 40$^{th}$ seconds in a 1 min recording. The top right plot shows the relative location of individual animals across recoding time. The bottom right plot shows the average speed of animals. When light is turned on or off, animals show substantial acceleration. 5x playback.

**Supplementary Video 4.** Video shows animals' natural swimming in 3% ethanol condition. The left panel was recorded between 0 ~ 1min and the right one was recorded between 4~5min. Each color represents an individual's trajectory. 5x playback.
